# Impact of the species compartment definition on quantitative modeling of microbial communities

**DOI:** 10.1101/018010

**Authors:** Marko Budinich, Jérémie Bourdon, Damien Eveillard

**Affiliations:** LINA UMR 6241, Université de Nantes, EMN, CNRS, France

## Abstract

Recent advances in genome-scale metabolic network reconstruction paved the way to the use of quantitative modelings such as FBA. However, despite the great interest of these techniques to tackle quantitative features, microbial community modeling remains unclear. Whereas studies represent a microbial community with several compartments for each microbial strains and their common pool, others advocate for the use of a single compartment that combines all reactions. Here we show that both modelings lead to different optimal quantitative solutions. This study illustrates this difference by the use of the flux module technique, that describes, in a compact way, the optimal solution space as computed by FBA-like techniques. For application, this paper computes the flux modules of a hot spring microbial community (represented by *Synechococcus spp., Chloroflexus and Roseiflexus spp.*) and a microbial methagenic system (*Desulvovibrio vulgaris* and *Methanococcus maripaludis*) sulfate reducing bacteria), while emphasizing the quantitative changes that occurs when one assumes either the consortium as a “single compartment” or a multiple compartment.

## Introduction

Following the surge of high throughput experiments to investigate microbial ecosystems, network systems ecology techniques risen their interest by proposing a qualitative description of ecosystems. For instance, by focusing on “who is there and who is not” [13], these techniques emphasize networks of microbes that co-occurred. However, the functional understanding of these communities remains often challenging when techniques solely consider qualitative community description rather than introducing (partial) quantitative knowledge. Considering the metabolic network of a community is one way to overcome this weakness [21]. One assumes the microbial ecosystem behaviors driven by the metabolic reactions encoded by each microbial genomes. In other words, thanks to multiple metabolic enzymes, microbes interplay within the environment and promote a whole metabolic network responsible for quantitative behavior [6, 3]. Once the metabolic network identified, constraint-based modelings are standardly used to reproduce quantitative properties of the whole metabolic network at a molecular resolution (see [11] or [18] for review)

However, whereas these modeling techniques such as FBA-like approaches became standard modeling routines for single cell systems, their applicative conditions remain to be deeper investigated when applied on microbial communities. In particular, microbial community metabolic network could be either reconstructed as a single integrated network by merging all reactions monitored by all bacterial strains (i.e. Single Cell Hypothesis - SCH), or considering natural boundaries between species by considering strain specific genes (i.e. Multiple Compartment Hypothesis - MCH) as already promoted in tissu specific networks [2, 10] or well-studied human gut microbial community [14]. Because both assumptions implies a distinct experimental efforts, this study proposes an analysis of SCH and MCH consequences on microbial community metabolic models. It is worth to notice herein that such an approach is a natural extension of a previous work of [9]. In 2010, Niels Klitgord and Daniel Segrè proposed a study of the impact of compartmentalization in metabolic flux models via the consideration of organelles within the metabolic network of yeast. However, at the time of the study, no genome-scale metabolic description of microbial community was available. We propose to overcome this weakness by taking benefits from recent biotechnological progresses. We complete the previous study by exploring herein the difference of both - MCH and SCH - hypotheses on two distinct genome-scale community metabolic networks. First, we analyze a microbial mat model that uses three microbial strains: Cyanobacteria, filamentous anoxygenic phototrophs and sulfate reducing bacterias [17], to analyze quantitative and qualitative differences in the predictions; secondly, we use *Desulfovibrio vulgaris* and *Methanoccocus maripaludis* microbial system, a classical example of syntrophic growth in anaerobic environment [16].

## Results

For the sake of application, one first considered the phototrophic microbial community system during day light composed *Synechococcus spp.*, called SYN, filamentous anoxygenic phototrophs related to *Chloroflexus* and *Roseiflexus spp.*, called FAP, and sulfate reducing bacteria (SRB) [17]. This community consumes CO_2_ and releases O_2_ by photosynthesis. As a byproduct of the rubisco activity, glycolate is produced by SYN, which will be later used as an organic substrate by FAP, along with acetate. Besides, SRB can consume organic compounds and reduce sulfate using H_2_. MCH community model describes a metabolic network for each strain as well as external metabolites such as H_2_, O_2_, NH_3_, glycogen and acetate (136 reactions) [17]. As a modeling contribution, a community biomass function was included to represent the ecosystem growth plus one extra reaction for preserving O_2_ / CO_2_ ratio as used by rubisco. As reported in [17], the so-called “Pool model” represents the SCH community model (59 reactions: 48 core and 11 exchange reactions) - see Figure 2 and Supp for details.

Both MCH and SCH models reproduce previous results and are qualitatively consistant with available experiments [16, 22] (see Figure 1). Both models show similar range of biomass increase when light increase. Naively, these similar predictions may lead to over-interpret that both model are identical, which do not advocate for the use of MCH that is experimentally expensive. However, solely, these interpretations are not sufficient and might lead to misunderstandings. To overcome these shortcomings, one must indeed consider extensive simulations to investigate all solutions as provided by both SCH and MCH models, and not a unique optimal solution. Indeed, while constraint-based modelings describe sets of fluxes that go through all metabolic reactions at equilibrium, above simulations represent, among all fluxes, a unique flux combination that maximizes one given objective - herein the biomass production. More recent Flux Variability Analysis (FVA) improves this unique description by pinpointing all satisfying solutions over a wide range of environmental conditions. Promoting a systematic exploration of these solutions, Flux Module (FM) technique [12] analyzes how, among all solutions, some reactions are systematically correlated - emphasizing subnetworks that connect a subset of substrates and products [7]. These subnetworks or modules are unique and result from all potential quantitative solutions. Implicitly, different modules imply different quantitative predictions (mathematically called optimal solution space). From a biological viewpoint, its application on single cell organisms shows modules as a sensitive description of biological functions (see supplementary for *E. coli* modules).

**Figure 1:**
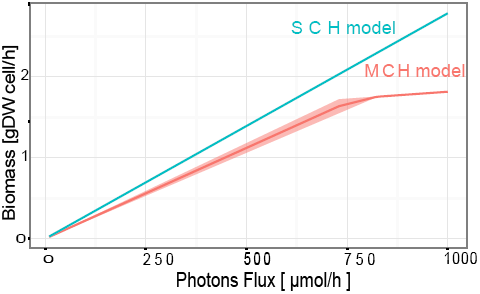
Quantitative simulations of Multiple Compartments Hypothesis (MCH) and Single Compartments Hypothesis (SCH) models of a hot spring microbial mat system. Flux variability simulation results show an increase of the whole community biomass under a range of light conditions. Simulations of MCH and SCH metabolic models are pictured in red and blue respectively.

MCH and SCH models produce distinct modules, which clearly emphasizes fundamental differences between MCH and SCH solutions. SCH shows only one module (purple in Figure 2), containing 31 reactions (52.2% of overall reactions): 20 reactions covered by MCH modules and 11 not previously highlighted. 7 MCH module reactions do not belong to the SCH module. MCH SYN reactions are decoupled from other networks, confirming previous studies [17] that highlights SYN as a primary producer for all possible microbial interactions. Complementary, FAP and SRB are linked via acetate and H_2_ metabolisms. As additional differences, the first glycolysis phase (R1-R2) and pentose phosphate reactions (R5-R9) are connected in SCH, which is not true when each organism is considered separately. SCH module is independent from uptake reactions; whereas MHC modules depict acetate processing of FAP and SRB linked to O_2_, H_2_ and CO_2_ exchanges.

**Figure 2:**
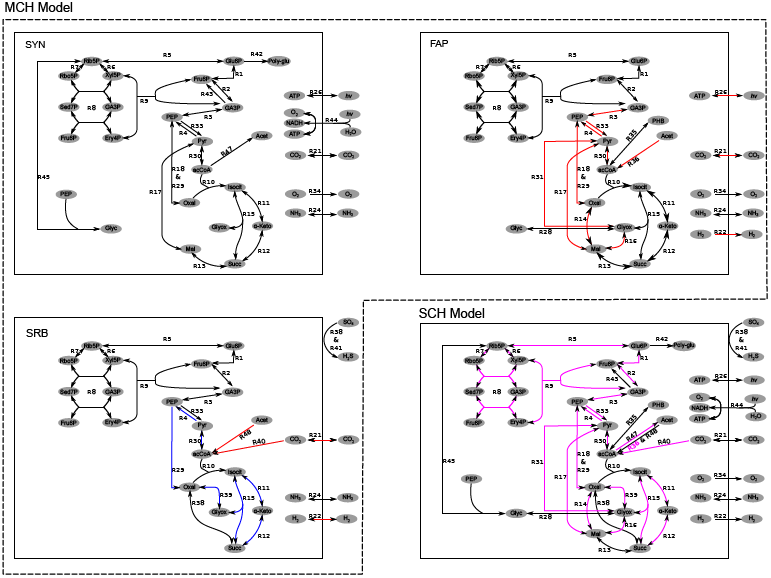
Description of metabolic networks related to the microbial mat community system and corresponding modules illustrations. SYN, FAP and SRB depict bacterial strains of the MCH metabolic model. SCH model represents the same metabolic system with no consideration of the compartments while conserving the naming convention of the MCH networks. For the sake of illustration exchange reactions between compartments are not shown. MCH model reveals 2 modules (26.5% of the whole set of metabolic reactions). One module contains 28 reactions (red) that span through FAP and SRB, whereas another (blue) involves 8 reactions. Reactions of the SCH module are depicted in purple.

For the sake of generalization, a similar modeling comparison was applied on a methanogenic microbial system composed of *Desulfovibrio vulgaris* and *Methanococcus maripaludis*. *D. vulgaris* uses lactate fermentation and sulfur reduction to gain energy, while producing gaseous hydrogen. *M. maripaludis* uses hydrogen to reduce CO_2_ into methane, which avoid the accumulation of H_2_ that might decrease the chemical energetic potential of *D. vulgaris*. The corresponding MCH model owns 243 reactions (respectively 145 and 97 reactions for *D. vulgaris* and *M. maripaludis*) and the SCH model is composed of 221 unique reactions, after deletion of redundant reactions. Again MCH and SCH modules are different (see Supplementary). Both models show a unique module: 124 reactions (48.6% of MCH model) and 187 reactions (84.6% of SCH model). Reactions related to H_2_ and acetate transport are not related in SCH model, whereas they are in MCH. Additionally, pentose phosphate cycle reactions of *D. vulgaris* and *M. maripaludis* are linked in SCH module but not in MCH, which might lead to misunderstand bacterial interpretations.

## Discussions

Despite similar quantitative simulations, this study shows significant differences between SCH and MCH. However, this communication do not advocate for either of both modeling assumptions. Biologically, SCH models have been widely employed to study metabolite exchanges between species (e.g., cocultures [20][5] or species within a complex environment [8]), whereas MCH models have been used to describe microbial communities, where each member seeks to maximize their own biomass [19]. Both assumptions are equivalent when one is interested by predicting overall quantitative behaviors of a microbial community, which is mostly explained by similar exchange reactions between SCH and MCH models. A protocol driven by SCH might be mostly sufficient for overall predictions with non further functional investigations. Reversely, MCH driven protocols present a significant cost to decipher boundaries between species and origin of genes within a meta-genome [18], but appear as necessary to investigate fine quantitative interactions within the community.

From a methodological viewpoint, this study advocates for the use of Flux Modules to compare metabolic models. Modules represent an abstraction of all Flux Variability simulations for a given metabolic model. Indeed Flux Module technique is a natural way to resume the methodological work of [9] that proposes an extensive analysis of yeast metabolic flux estimation with and without compartmentalization. Since our study pinpoints similar conclusions to [9], both studies reinforces the need for further constraint-based modelings dedicated to multiple compartments simulations as motivated by [22, 23].

## Acknowledgments

M.B.is supported by CNRS grant, M.B., J.B. and D.E. are supported by GRIOTE project. D.E. is supported by ANR grant SAMOSA.

## 1 Supplementary materials

### 1.1 Flux Modules are biologically relevant: illustration on modules for *E. coli* metabolic network in aerobic and anaerobic growth conditions

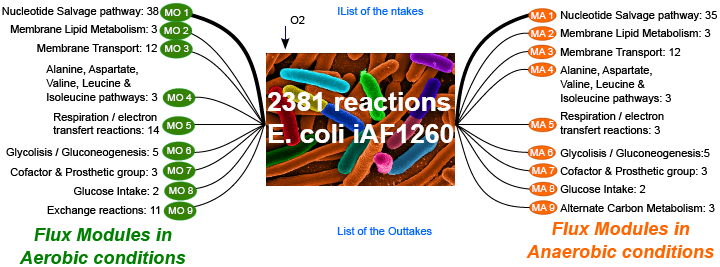

Above figure depicts modules of *E. coli* in both aerobic and anaerobic growth conditions. The metabolic model composed of 2381 reactions is analyzed in aerobic (left) and anaerobic conditions (right). For each conditions, one extracts 9 modules that are composed of distinct numbers of reactions (line width is proportional). Each module is associated to the pathways in which the flux module reactions are involved. Modules are in accordance to biological conditions. When challenged by an oxidative stress, most of flux modules of *E. coli* are conserved, except exchange reaction, respiration & electron transfert, alternate carbon metabolism, which is in accordance to physiological knowledge. To a lesser extent, nucleotide salvage pathway is impacted by oxygen growth conditions.

### 1.2 Metabolic modules of a methanogenic microbial system composed of *Desulfovibrio vulgaris* and *Methanococcus maripaludis*

The metabolic model of *D. vulgaris* contains 145 reactions [22], whereas *M. maripaludis* model is composed of 97 reactions [16]. In order to link both strains within a MCH model, we duplicated exchange reactions of H_2_ in order to import/export the metabolite with either the other microorganism or environment. A similar procedure was done for Formate, Acetate and CO_2_, which overall introduces 12 exchange reactions. Finally, the whole ecosystem biomass was design to fit biomass functions of *D. vulgaris* and *M. maripaludis* as already published, while maintaining a respective proportion of 2:1 for both strains. SCH model of this community consist in merging both metabolic networks and removing all replicated reactions for considering one unique representative reaction. As results, SCH model is composed of 221 unique reactions, as a reduction of 243 reactions MCH model (respectively 145 and 97 reactions for *D. vulgaris* and *M. maripaludis*).

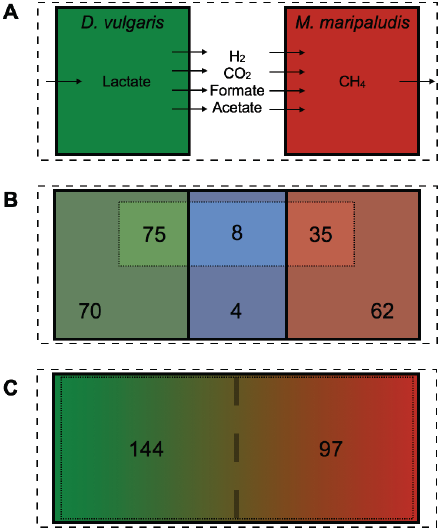

The SCH presents a disadvantage regarding the interplay between interchange fluxes; in this case, acetate and H_2_ are major players of electron transfer in anaerobic systems which role is an active area of research. Besides, for these two systems, the SCH links the fluxes of pentose phosphate system which could impact interpretations in future *in silico* developments

